# Computational Prediction of Synthetic Circuit Function Across Growth Conditions

**DOI:** 10.1101/2022.06.13.495701

**Authors:** Breschine Cummins, Robert C. Moseley, Anastasia Deckard, Mark Weston, George Zheng, Daniel Bryce, Joshua Nowak, Marcio Gameiro, Tomas Gedeon, Konstantin Mischaikow, Jacob Beal, Tessa Johnson, Matthew Vaughn, Niall I. Gaffney, Shweta Gopaulakrishnan, Joshua Urrutia, Robert P. Goldman, Bryan Bartley, Tramy T. Nguyen, Nicholas Roehner, Tom Mitchell, Justin D. Vrana, Katie J. Clowers, Narendra Maheshri, Diveena Becker, Ekaterina Mikhalev, Vanessa Biggers, Trissha R. Higa, Lorraine A. Mosqueda, Steven B. Haase

**Affiliations:** Montana State University, Bozeman, MT, U.S.A.; Duke University, Durham, NC, U.S.A.; Geometric Data Analytics, Inc., Durham, NC, U.S.A.; Netrias LLC, Annapolis, MD, U.S.A.; SIFT, LLC, Minneapolis, MN, U.S.A.; Strateos, Inc., Menlo Park, CA, U.S.A.; Rutgers University, New Brunswick, NJ, U.S.A.; Raytheon BBN, Cambridge, MA, U.S.A.; Texas Advanced Computing Center, Austin, TX, U.S.A.; UW Biofab, Seattle, WA, U.S.A.; Ginkgo Bioworks, Boston, MA, U.S.A.

**Keywords:** Robustness, Circuit, Prediction, Performance, Topology

## Abstract

A challenge in the design and construction of synthetic genetic circuits is that they will operate within biological systems that have noisy and changing parameter regimes that are largely unmeasurable. The outcome is that these circuits do not operate within design specifications or have a narrow operational envelope in which they can function. This behavior is often observed as a lack of reproducibility in function from day to day or lab to lab. Moreover, this narrow range of operating conditions does not promote reproducible circuit function in deployments where environmental conditions for the chassis are changing, as environmental changes can affect the parameter space in which the circuit is operating. Here we describe a computational method for assessing the robustness of circuit function across broad parameter regions. Previously designed circuits are assessed by this computational method and then circuit performance is measured across multiple growth conditions in budding yeast. The computational predictions are correlated with experimental findings, suggesting that the approach has predictive value for assessing the robustness of a circuit design.

## Introduction

Synthetic biologists often seek to construct synthetic genetic circuits that accept an input signal(s) and, in response, produce some output signal. An example would be biosensors with environmental or medical applications [1-3]. Synthetic genetic circuits have been constructed with network engineering principles that incorporate non-linear interactions within the circuit. Non-linearities are inherent in genetic circuits at multiple levels including, for example, cooperativity in protein/DNA binding and protein-protein interactions.

Although network function can be modeled by a system of differential equations, it is largely intractable to measure the parameters that determine circuit behavior within a cell.

Moreover, those parameters are likely to be noisy and will certainly change as the cell states change due to normal growth and division or as cells encounter changing environmental conditions (e.g. temperature, nutrient availability). Parameters might also be affected by genetic differences between species or between distinct strains of the same species. Thus, circuit performance might not be reproducible unless genetics and growth and environmental conditions are tightly maintained. Not surprisingly, the quest for reproducibility has relied heavily on methods for precisely defining and controlling experimental conditions across laboratories [4]. In some cases, even the best efforts to control experimental conditions do not yield reproducible results [5].

The difficulty in maintaining experimental conditions within a laboratory highlights a fundamental challenge for synthetic biology - if reproducible results cannot be easily achieved in laboratory conditions, how will circuits perform when deployed into the environment or within the changing environments of an organism?

An alternative to achieving reproducibility by controlling experimental conditions is to design circuits that perform the desired functions across a broad range of parameter regimes. The function of such circuits should be robust to noisy or changing parameters, and in turn, to environmental changes that might be encountered during a deployment. The basis for this approach is the realization that robustness should have a positive effect on reproducibility. The challenge of this approach is to design robust circuits without substantial information about the parameter regimes in which they will operate. Approaches have been developed to establish input/output (dose/response) relationships for particular interactions within circuits[6].These measurements can be expensive and time consuming. At the same time, these relationships only hold for the experimental conditions in which they were interrogated, so they are not generalizable to other conditions. New design approaches that combine parts-level approaches with circuit scoring methods appear to increase robustness in designs [7].

Here we describe an alternative approach for generating robust circuits that will perform reproducibly across growth conditions. The approach enables synthetic biologists to computationally assess the function of a particular circuit design broadly across parameter space. We first make computational predictions about the function of previously designed circuits [8] across parameter space, and then test circuit functions across multiple growth conditions to assess the accuracy of the computational predictions.

## Results

Predictions of the performance of various synthetic logic circuits were made using the Python 3 package Dynamic Signatures Generated by Regulatory Networks (DSGRN) [8, 9]. This tool allows the comprehensive modeling of the dynamical behavior of a network topology over parameter space in a computationally scalable manner. Network topology means the interaction structure of anonymous gene products; specific builds of a network are not explicitly modeled, although implicitly they are assumed to correspond to some parameterization of the network topology. The dynamical behavior being modeled includes, but is not limited to, truth tables of logic functions, which is the dynamical behavior considered in this work. The ability of DSGRN to comprehensively predict behavior over all of parameter space permits the assignment of a robustness score as a percentage of parameters at which the desired logical behavior is observed. The robustness score provides a relative measure of the ability of a network topology to reproduce a specific observed behavior among a collection of networks.

Recently, a set of functional synthetic two-input logic circuits (AND, OR, NAND, NOR, XOR, XNOR) built in the yeast *S. cerevisiae* were published [10]. These logic circuits were built out of more primitive parts: CRISPR-constructed NOR gates where the regulatory molecule is a short segment of RNA called a guide RNA (gRNA). NOR logic was chosen because every logical expression can be constructed from an arrangement of NOR and NOT gates. The inclusion of CRISPR reduced transcriptional leakage under repression. Each logic function was composed of four strains, one for each Boolean input state (00, 01, 10, 11), where the expression of the input gRNA was controlled by the presence or absence of the associated promoter. Various combinations of the CRISPR NOR gates were constructed for each logic circuit. It was found that certain builds of each logic function exhibited clean and reproducible performance; however, there was variation across the builds in functional performance within each logic function. In the context of the study, the terms functional and performance refer to a circuit’s ability to express fluorescence (ON) or not (OFF) as intended by the circuit’s logic, or truth table, given a Boolean input state.

Table 1 shows DSGRN predictions of the relative performance of these six circuit topologies across experimental condition space conditioned on the fact that the cells grow. Theoretically, the robustness score can vary between 0 and 1, see [11] or the computation and [12] for a similar method. In reality, it is extremely unlikely if not impossible for a robustness score of 1 to be attained, but complex circuits can attain robustness scores of 0.5 or higher. It is important that these numbers are not interpreted as probabilities; only the rank order of the circuits is meaningful. DSGRN predicts that NOR should be the highest performing circuit across media conditions, followed by OR. Next, AND is expected to perform a little better than NAND, and lastly XOR is expected to perform a little better than XNOR. From previous experience it is known that these scores are small in absolute value compared to what is seen for more complex networks, and therefore these rankings are not well-separated. This lack of separation in scores is due to the simplicity of the circuits that do not have functionally redundant motifs. An evaluation of these circuit predictions in comparison to data follows.

**Table 1.** DSGRN predictions for relative ordering of synthetic logic circuits.

| Circuit | NOR | OR | AND | NAND | XOR | XNOR |
| --- | --- | --- | --- | --- | --- | --- |
| DSGRN Robustness Score | 0.077 | 0.067 | 0.059 | 0.053 | 0.034 | 0.026 |

As published previously in [13], a number of experiments were performed on the CRISPR circuits [10] under various growth conditions. Only one build of each of the six logic functions was studied for performance across media conditions in the set of experiments analyzed here. Each circuit’s ON state results in the expression of green fluorescent protein (GFP) so circuit output can be assessed by the fluorescent intensity of a cell. Both plate reader and flow cytometry data were collected over five experimental rounds. Growth conditions, representing rich media, minimal media, high osmolarity, poor carbon sources, and high temperature were varied by using four different media respectively (YEP 2% Dextrose, Synthetic Complete, Synthetic Complete containing 1% Sorbitol, and Synthetic Complete containing 2% Glycerol and 2% Ethanol) and two temperatures (30°C and 37°C). As these growth conditions can slow the rate at which yeast grow, a time-series protocol was employed to account for variable growth rates across conditions. Two of the five rounds did not exhibit sufficient cell density for reliable analysis or comparison to predictions and were excluded from the analysis below. Sufficient cell density was taken as a proxy for cell growth and was set at a threshold of 1,000,000 cells/mL. In addition, the Ethanol media condition frequently did not exhibit sufficient cell density and was excluded from much, though not all, of the analyses below.

Flow cytometry data in arbitrary units are aggregated over these experiments and shown in Figure 1. The plots show the distribution of geometric means of flow cytometry samples per strain excluding the Ethanol media condition. The x-axis and legend colors together describe the truth table for each circuit. Considering NAND in the lower left for example, the input states 00, 10, and 01 are all intended to be ON and 11 is intended to be OFF. The figure shows that in general, NAND performs as intended since the geometric means of the flow cytometry distributions are higher in the intended ON state compared to the intended OFF state. Some circuits do not have a good separation in geometric mean between strains intended to be ON and those intended to be OFF, e.g. AND and XNOR, but there are two clear examples of unexpected output. One is the NOR 11 strain, which expresses fluorescence when it should not, and OR 10, which does not express fluorescence when it should. Using several complementary analysis techniques with various strengths, we quantify these qualitative observations.

**Figure 1.**
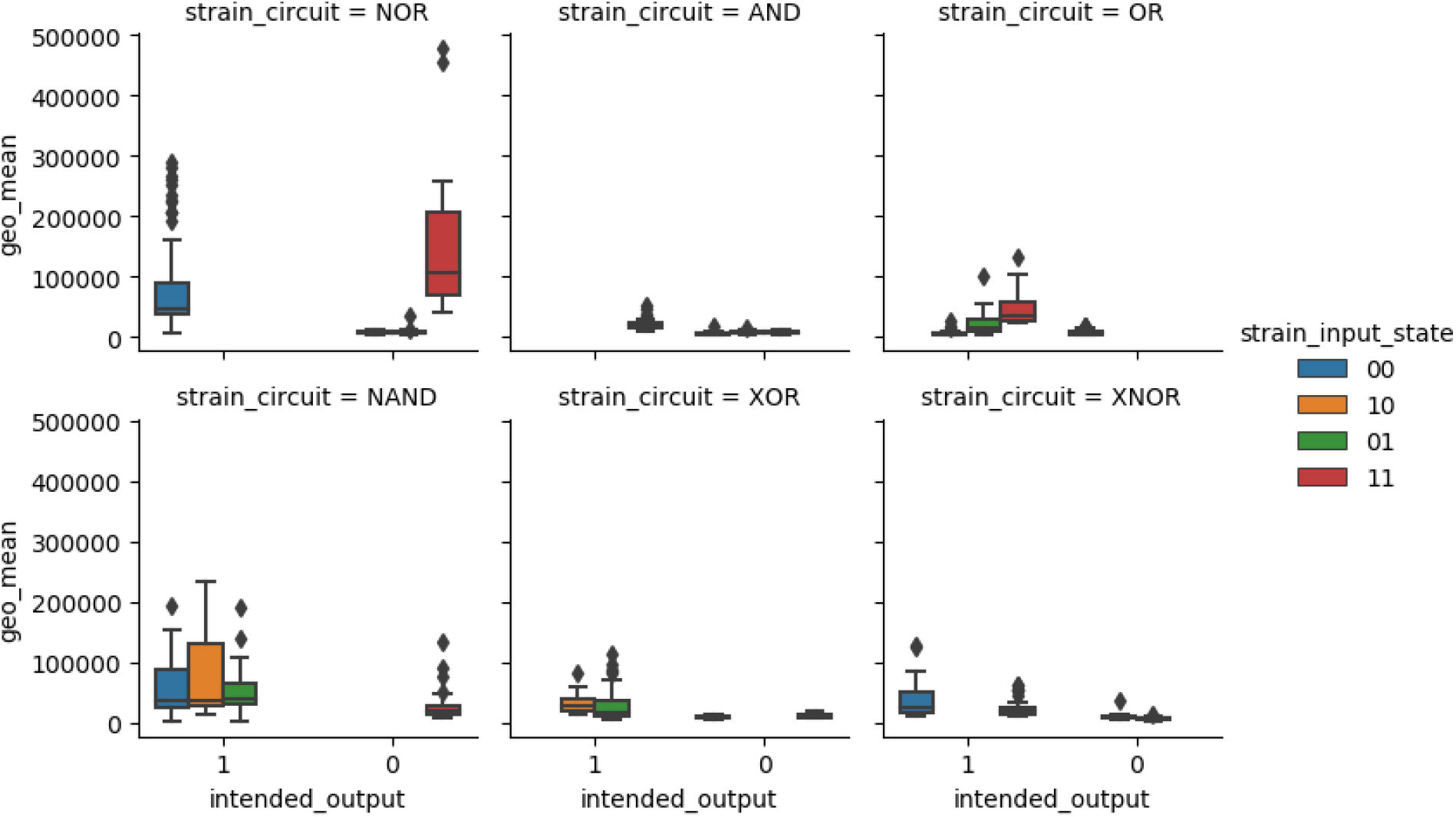
Data from Round 3 [13] for each of the 6 circuits. The Ethanol media condition is not included due to generally poor growth. The x-axis shows the expected state to be achieved by the circuit. Intended output 1 corresponds to ON, or high fluorescence, and intended output 0 corresponds to OFF, or low fluorescence. The colors appear consistently in the same order, so, for example, in the top middle panel for AND where the colors are not visible, the 11 input condition is associated with intended output 1 and the three other inputs are associated with intended output 0.

Figure 2 shows the quantification of circuit performance over three rounds based on flow cytometry data, this time including the Ethanol media condition. The technique used to quantitate circuit performance is called Wasserstein cut scoring (see Methods), which ranks between 1 and 9 the similarity of the data to each of the sixteen logical truth tables that are possible with two inputs. This is done by taking pairwise distances between the flow cytometry histograms and using the control histograms plus a graph-based method to provide a numerical score to each possible logical truth table. These scores are not very meaningful on their own, so we choose to use rank as a better indicator of relative circuit performance. The top-ranked truth table for each set of histograms associated with input conditions 00, 01, 10, and 11 for a circuit and an experimental condition is considered to be the best fit to the data.

**Figure 2.**
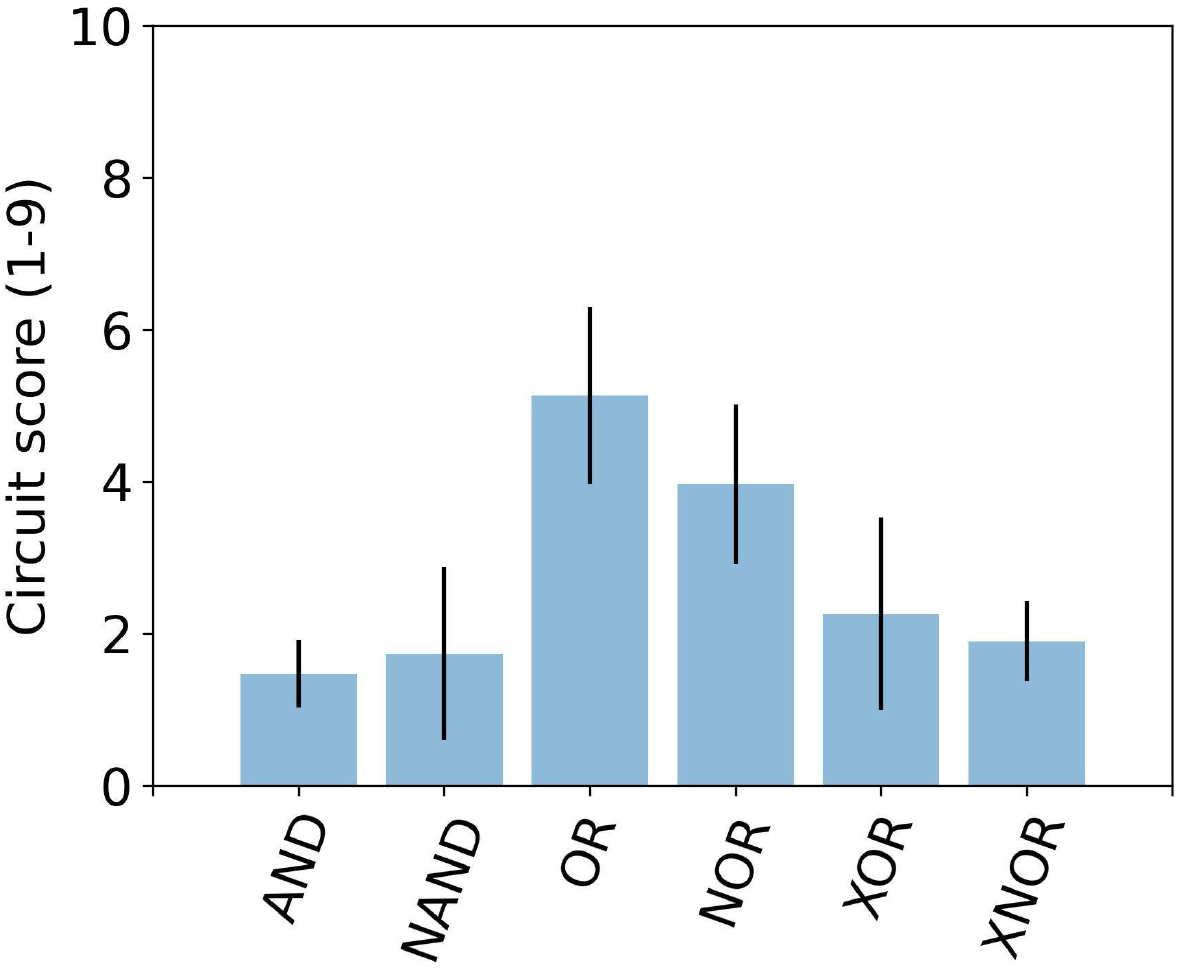
Circuit performance across media conditions. The mean and standard deviation (using Python’s numpy.std) of the Wasserstein cut score (see Methods) for each synthetic logic circuit across all three rounds, including the Ethanol media condition. Lower ranks indicate better performance. This figure is very similar to Figure 6 in [13], but is generated from raw log10-transformed data instead of ETL processed data.

The mean and standard deviation of the rank of the *expected* truth table is shown for each circuit in Figure 2, so that lower values indicate better performance. For example, the data of the AND circuit most closely represented the AND truth table by ranking in first or second place on average, resulting in the best score over all circuits. On the other hand, the OR circuit data matched the OR truth table only at rank 5 on average. In general, the data for the OR circuit resembled other logical truth tables, resulting in the worst score over all circuits. The analysis in Figure 2 does not show significant differences between circuits. However, the averages indicate worse performance by OR and NOR compared to the other four circuits. On average, OR and NOR had the correct truth table ranked 4-5 out of 9 places, despite being predicted to be the top performers by DSGRN.

The scores shown in Figure 2 are computed by aggregating information from all four strains that compose a circuit for an experimental condition and time point. The circuit strains showed varying growth and density across experimental conditions and time points, and not every circuit had all four strains with sufficient cell density and event sampling to be confident of analysis in every condition at every time point. Therefore, the variance in Figure 2 is generated from differing numbers of experimental conditions and time points that vary across circuits, which is summarized in Table 2. The proportion of experimental conditions score (PEC) is the proportion of experimental conditions used to compute the scores shown in Figure 2 out of the total number of experimental conditions per circuit in which at least one strain of the circuit had sufficient cell density. The PEC score varies between 0 and 1 and is a different measure of robustness of circuit performance. For example, AND has a better Wasserstein cut score than XOR in Figure 2, but XOR was scored across a larger number of conditions, which may explain its higher variance.

**Table 2:** Proportion of experimental conditions with sufficient cell density (PEC) for Wasserstein cut score analysis.

| Data Scores | NOR | OR | AND | NAND | XOR | XNOR |
| --- | --- | --- | --- | --- | --- | --- |
| PEC | 0.88 | 0.83 | 0.71 | 0.78 | 0.91 | 0.78 |

We examined the circuit performance scores across different media conditions to ask whether any particular growth condition was biasing the scores (Figure 3). Results are shown only for the round with the highest growth rates and cell densities, so there are even fewer data points per bar than in Figure 2. No single growth condition appears to bias results across different truth tables.

**Figure 3:**
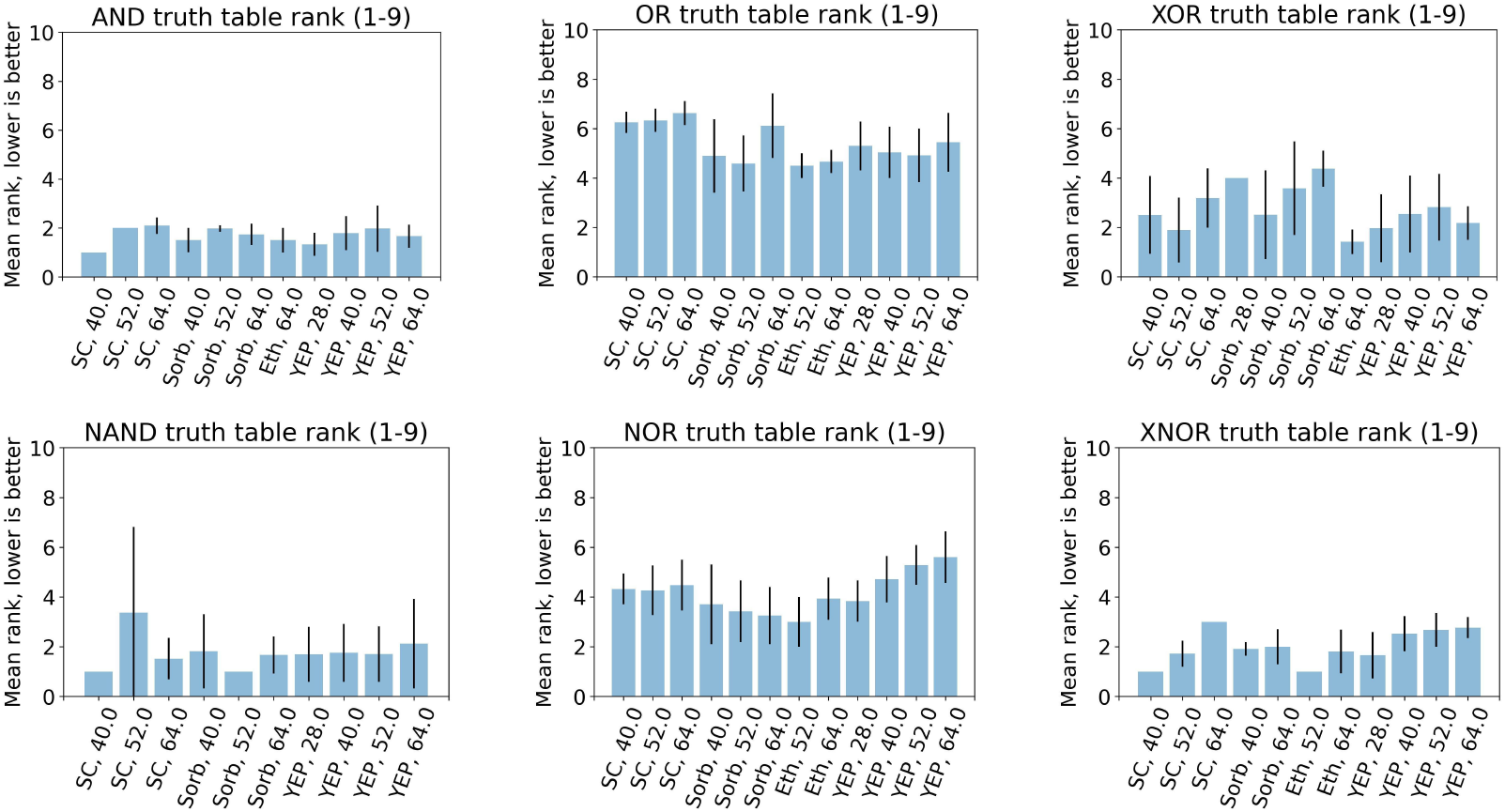
Circuit performance separated across media and time. The mean and standard deviation of the Wasserstein cut score for each synthetic logic circuit. When error bars are absent, this indicates a standard deviation of zero. The categories on the x-axis are media and time point (hr) and the temperature is fixed at 30°C. Notice that the performance of each circuit is over a slightly different set of conditions. The missing conditions are those in which there was insufficient cell density to have confident results. The data shown are from Round 3 [13] that exhibited high cell densities consistently.

The Wasserstein cut score does not account for fold change in fluorescence between ON and OFF circuit states. In particular, the histograms of ON and OFF states may even have a fold change near 1. Having a high fold change with a low variance is a proxy for a confidence level in the Wasserstein cut score. Results from two analyses that examine fold change in fluorescence are shown in Figure 4. The top panel shows a boxplot for each circuit, where each bar summarizes the distribution of the geometric mean for flow cytometry data. The orange bars correspond to the strains that should produce GFP and the blue bars correspond to those that should not. For example, the NOR circuit shown in the top row has three strains associated with the blue bar, corresponding to logical input states 01, 10, and 11, while the orange bar is associated with samples of the single strain NOR 00. This panel shows that while the NOR strains have a good separation in median between ON and OFF states, there is substantial overlap in their distributions. The OR strains, another poor performer according to the Wasserstein cut score, have the second greatest overlap. XNOR and AND (the best performer according to Wasserstein cut scoring) have the least overlap between ON and OFF distributions; however, their medians are not well-separated.

**Figure 4:**
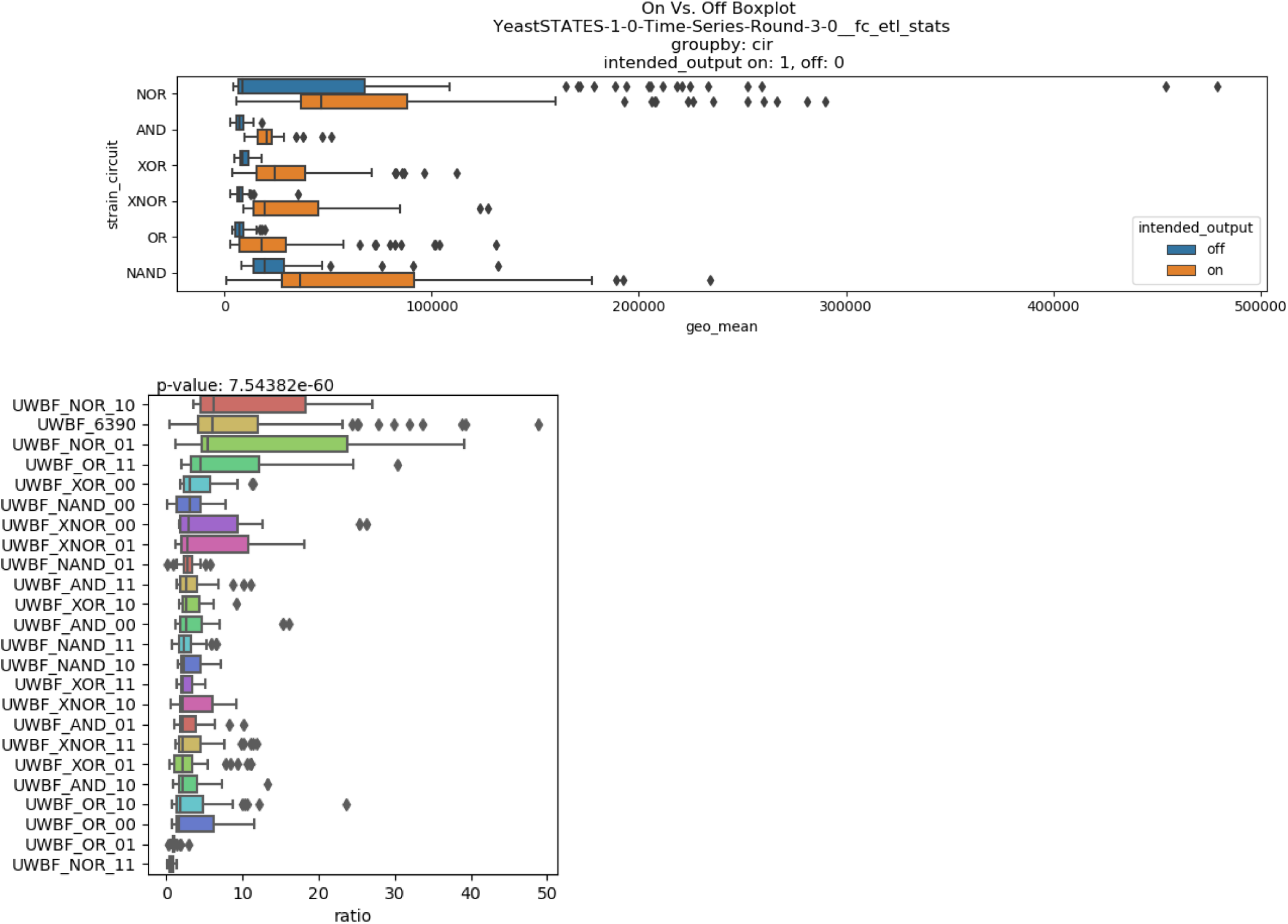
Circuit performance using the Performance Metrics (top) and Data Diagnosis (bottom) analysis packages [11, 13-15] on flow cytometry data. The strain labeled UWBF_6390 in the bottom panel corresponds to the NOR 00 strain. The bottom panel is very similar to Figure 7 in [13] but is generated from data in arbitrary units instead of ETL processed data.

The bottom panel of Figure 4 shows the fold change expressed as ON median/OFF median ratio for each of the four strains per circuit independently. High ratios are indicative of a high fold change. The top row contains the strain with the best fold change in median, which is NOR 10, and the bottom row shows the worst, which is NOR 11. Each of these two strains is not supposed to produce GFP, so the medians of their flow cytometry histograms form the denominator in the ON/OFF ratio. The numerator is the median of the strain of the NOR circuit that is supposed to produce GFP, i.e. NOR 00 (labeled UWBF_6390 in the figure). It is seen that NOR 11 and OR 01 are particularly poor performers, since their fold change is near 1 with low variance, indicating that it is difficult to distinguish the expected ON and OFF flow cytometry histograms.

Two hypotheses for the underperformance of OR and NOR are possible. 1) DSGRN predictions are incorrect and the gates lack any kind of robustness or 2) the strains are not representative of OR and NOR gates due to incorrect build, human error, or circuit malfunction. With further examination, it was observed that the provenance of the OR strains is not clear, and that there may have been an error in the labeling or construction of one or more OR strains. This is backed by inconsistencies between the reported strain sequence, the sequence uploaded to SynBioHub [11], DNAseq [16], and small RNAseq experiments.

Paired-end small RNAseq was performed to enable investigation of failure mechanisms of yeast gate circuits at the transcriptional level. Upon searching the raw sequencing data for the presence or absence of all gRNAs used to build the various circuits, disagreements were uncovered between the expected circuity and the observed circuity. Specific to the OR circuit strains, there were strains both missing expected gRNAs and containing unexpected gRNAs.

When these results were compared to DNAseq results and to annotations of the OR gate from the original publication [10] and in SynBioHub, disagreements were found between all four sources of information; see Figure 5 for the example of the OR 11 strain. This observation supports the hypothesis that either an unexpected build or mislabeling issue is responsible for the poor performance of the OR circuit. The NOR gate was not investigated.

**Figure 5.**
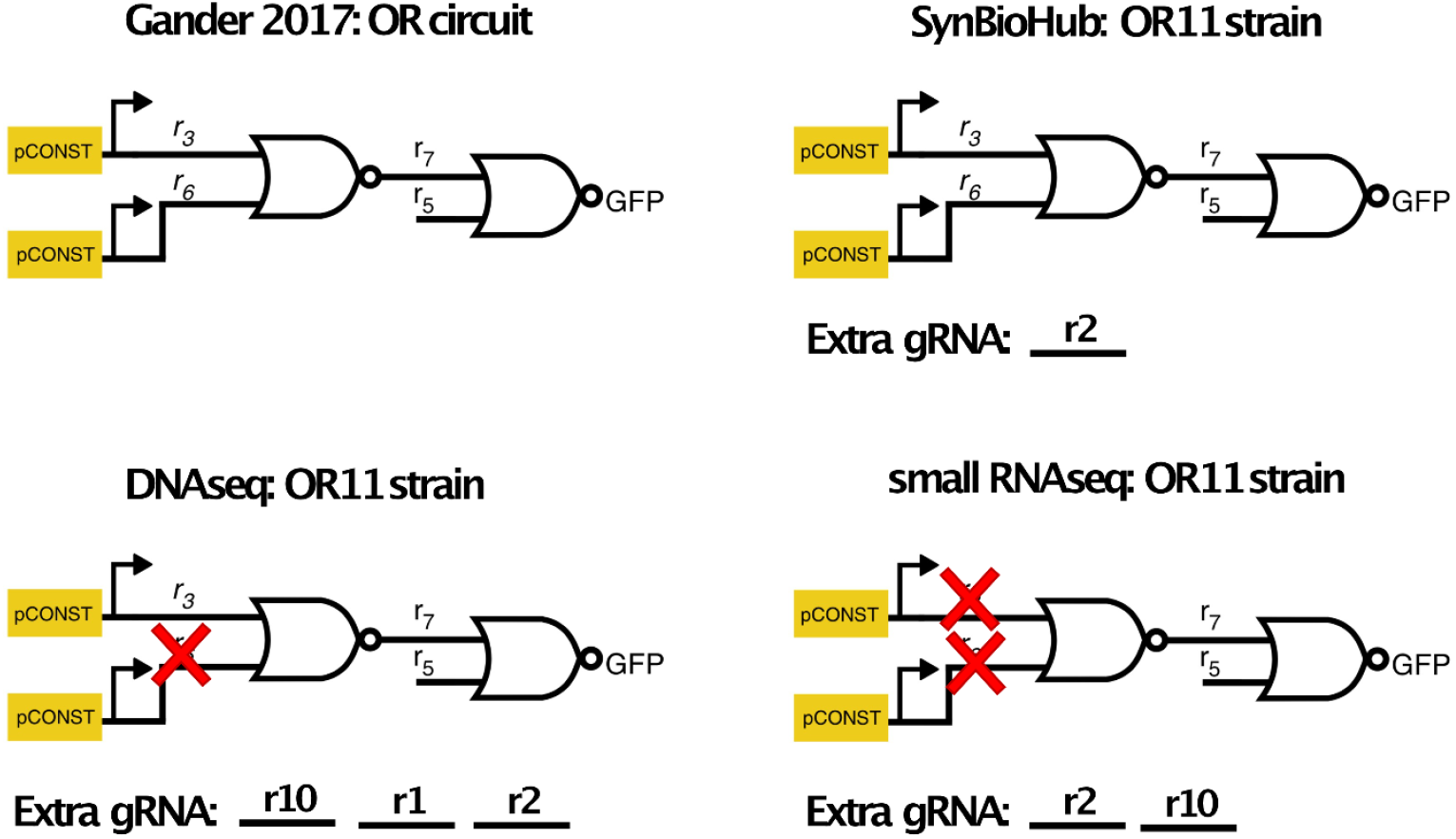
Expected and unexpected gRNAs for the yeast OR gate with an input state of 11 from various resources. Red “X” indicates the gRNA is missing from the resource.

## Discussion

In the absence of correct builds for OR and possibly NOR, DSGRN predicts the highest performing circuit to be AND with a close second to NAND (Table 1). It is seen in Figure 1 that the AND circuit has both the lowest mean and lowest standard deviation of the six circuits. AND performs slightly better than NAND, and both perform better than XOR and XNOR. Although this is not statistically significant, the trend is suggestive of predictive accuracy, particularly given the small numerical separation of the predictions. On the other hand, the predictions indicating that XOR should perform better than XNOR are not obtained. XNOR performs better than expected.

The discerning reader will have noticed that the DSGRN predictions in Table 1 follow in order of complexity, with the simplest circuit, NOR, ranked highest, and XNOR ranked lowest; see [10] for the full set of circuit topologies. It is not in general the case that DSGRN predictions track circuit complexity. The observation that more complex circuits may have greater robustness due to increased redundancy or other topological motifs that may increase robustness is also reported in [7, 17]. Thus, in the context of synthetic circuits there may be a natural tension between robustness and ease of circuit construction.

The experiments have shown that OR and NOR are performing poorly for reasons that are probably not due to a “natural” lack of circuit robustness. Ideally these circuits should be rebuilt and retested in the future, but the findings do indicate that circuit performance predictions might help identify strains where circuits do not reflect the expected design specifications. Moreover, the observation that DSGRN can predict the robustness of a particular circuit across multiple growth conditions suggests that this tool may be used to assess circuit robustness and performance at the design phase, before construction begins. The ability to computationally evaluate robustness in a highly scalable manner before circuit construction will likely provide a time and cost savings over evaluating circuit functions only after construction. Towards this end, we have implemented the DSGRN Design Interface, a user-friendly interface built around DSGRN that transforms DSGRN from a predictive tool to a design tool for 2-input feed-forward logic circuits. This tool has been used to assess the trade-off between circuit complexity and robustness [11].

Our findings indicate the utility of DSGRN to computationally evaluate the robustness of synthetic genetic circuit topologies, even before their construction. The results indicate that while the different parts utilized to construct these circuits likely influence the parameters that guide circuit performance, the topology itself is an important contributor to robust function. The ability to construct circuits that are robust across growth conditions has important practical benefits as any field deployment of a genetic circuit will likely encounter a broad range of changing growth conditions. Thus, reproducibility in circuit function can be improved by circuit design rather focusing only on maintaining tight control over experimental and growth conditions.

## Methods

### DSGRN

Dynamic Signatures Generated by Regulatory Networks (DSGRN) [8, 9, 18] is both a theoretical framework and software package for characterizing the long-term dynamical behavior of regulatory networks. It operates by assuming one parameter per node controlling decay and three parameters per edge controlling regulation. This high-dimensional parameter space is then rationally partitioned into a finite number of regions, where the partition is determined by identifying conserved dynamics. Any real-valued set of parameters in that region exhibits the same long-term behavior, such as oscillations and fixed points. A collection of 2 ^*n*^ fixed points can represent a logical truth table for *n* inputs; we investigate dynamical behaviors that represent logic with two inputs in this work. The decomposition of parameter space enables the comprehensive and rapid computation of all possible behaviors of a regulatory network, making it the ideal tool for predictions of robust behavior.

### Wasserstein cut score

The scores in Figure 2 are derived from the earth mover’s distance, or Wasserstein distance [19], between flow cytometry histograms. The pairwise distances between the histograms for each of the four input states (00, 01, 10, 11) lead to a graph-based score that permits a rank-ordering of the 16 possible truth tables for a two-input logical system. The 16 truth tables include the common AND, NAND, NOR, OR, XOR, and XNOR discussed in this manuscript, the constant truth tables ID and NOT ID, and also the less-used logic that is composed of the eight forms of IMPLY and NIMPLY. The code base is publicly available [20].

Let *h*_00_, *h*_01_, *h*_10_, and *h*_11_ be the flow cytometry histograms for each of the four input states and let *h*_*neg*_ and *h*_*pos*_ be the histograms for the negative and positive controls respectively. All possible combinations of the four histograms across technical and biological replicates for a single circuit, a single experimental condition, and a single time point are scored as described below. Let *d* represent the Wasserstein distance. Define the normalized Wasserstein distance *w* as

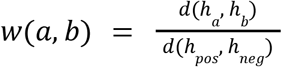

The normalized Wasserstein distance is a measure of how far apart two histograms are given the distance between the controls. It is used as a weight on the edges of the complete graph of four nodes representing the input states 00, 01, 10, and 11, see Figure 6.

**Figure 6:**
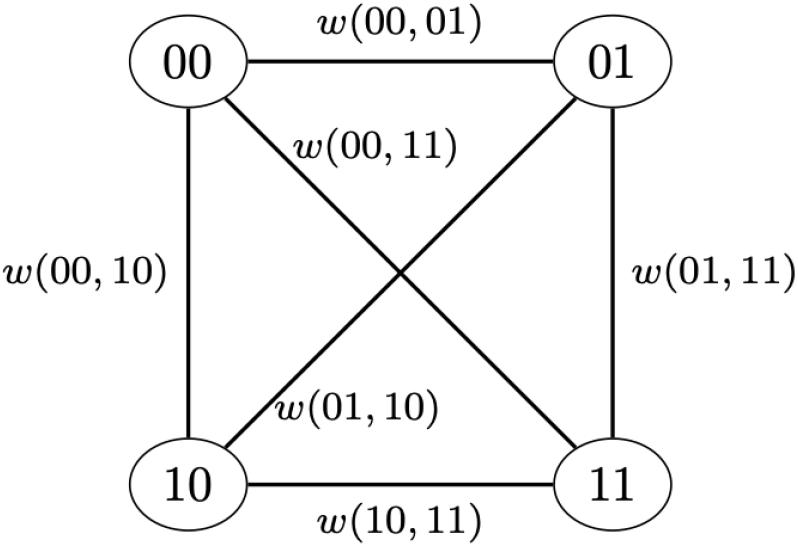
The Wasserstein cut scoring technique. The edges of the graph are weighted by the Wasserstein distance normalized by the distance between positive and negative controls.

We use the idea of a graph cut [21] to produce a score to rank-order the fit of truth tables to the data *h*_00_, *h*_01_, *h*_10_, and *h*_11_. We partition the nodes of the edge-weighted graph in Figure 6 to each of the seven possible two-partitions of the input states 00, 01, 10, and 11 (e.g. [00] and [01,10,11]; [00,11] and [01,10]; etc.). We then look at every cut-set (the set of edges joining nodes in one partition directly to nodes in the other partition) and take the average weight of the cut-set. A large average indicates that the histograms in the two partitions are far apart from each other. The largest such average cut score identifies the best partition of nodes for a truth table.

That partition corresponds to two truth tables, T and NOT T. For example, the partition [00] and [01,10,11] means that one set has an expected output value of 1 (ON) and the other has the expected value 0 (OFF). If 1 is assigned to the partition [00], then we have the NOR circuit, where fluorescence should be produced only if both inputs are absent. Likewise, if the partition [00} is assigned the state of 0, we have the OR circuit.

To decide between T and not T, the average is taken over the medians of the histograms in each partition. The input states associated with the higher average median are assigned 1, and the input states of the other partition are assigned 0. This truth table T is assigned the computed average cut score and the negation NOT T is given a score of infinity.

This results in a list of fourteen scored truth tables. The last two truth tables, the constant ones, are scored by a heuristic method that requires a hyperparameter (see code)[20]. One constant truth table is given a finite score and the other an infinite score. The absolute values of the scores themselves are somewhat fragile, so we choose to look at rank only, where the finite scores provide us with eight ranked truth tables and we assign rank 9 to every infinite negation. The top ranking truth table is considered to be the best fit to the flow cytometry data in terms of Boolean logic.

See [7] for another way of using the Wasserstein distance as a circuit robustness score.

### Flow cytometry data

See [13].

### gRNA quantification for yeast gate circuits from small RNAseq

FASTQ files from paired-end small RNAseq were searched for the presence or absence of gRNA sequences, specifically the sequence of the gRNA containing the target and handle sequences. Quality control was first performed on the small RNAseq data where quality scores were assessed using a sliding window of 4 nucleotides in length. Reads were dropped if the average quality score of a window was less than 15. This resulted in only ∼30% of both forward and reverse reads surviving quality control, ∼55% of only forward surviving and ∼2% of only reverse surviving. Therefore, only the forward reads were used to search for gRNA sequences. Additionally, gRNA abundances could only be compared within a sample, and not across samples as a normalization method was needed to make this possible. Small RNAseq reference beads were included in the experiments from Ginkgo, however, due to quality issues, the reference beads were not usable to normalize gRNA abundances. After quality control, the Levenshtein distance metric was used to identify reads in the fastq files that contained an 80% match to gRNA sequences used in the yeast gate circuits. The Levenshtein distance is a metric to measure how different two sequences of words are and was chosen to account for potential inaccuracies in sequencing.

## Data availability

Processed data and scripts used in the generation of figures in this manuscript are available in a public repository. The code packages cited in the manuscript are all open source.

## Author contributions statement

Conceptualization: BC, MG, TG, KM, SBH.

Data curation: BC, RCM, AD, D Bryce, TJ, MW, GZ, JN, KJC, JB, BB, TM, TTN, NR.

Formal analysis: BC, RCM, AD, TJ, MW, GZ, NR. Funding acquisition: D Bryce, MV, TG, KM, SBH. Investigation: JN, KJC, D Becker, EM, VB, TRH, LAM. Methodology: BC, RCM, AD, D Bryce, MG, JB.

Project administration: D Bryce, JB, MW, MV, SBH. Resources: JV, JN, NM, NG, JU, MV.

Software: BC, RCM, AD, D Bryce, TJ, MW, GZ, SG, JU, RPG, MG, JB, BB, TM, TTN, NR.

Supervision: D Bryce, JB, MW, MV, NM, TG, KM, SBH. Validation: AD, MW, GZ, JN.

Visualization: BC, RCM, AD.

Writing, original draft: BC, RCM, SBH.

Writing, review and editing: BC, SBH, MW, JB, TG, KM.

## Material Availability

N/A

## Acknowledgements

This document does not contain technology or technical data controlled under either U.S. International Traffic in Arms Regulation or U.S. Export Administration Regulations. Views, opinions, and/or findings expressed are those of the author(s) and should not be interpreted as representing the official views or policies of the Department of Defense or the U.S. Government. Approved for Public Release, Distribution Unlimited (DISTAR case 36346).

